# FabF and FadM Cooperate to Recycle Fatty Acids and Rescue Δ*plsX* Lethality in *Staphylococcus aureus*

**DOI:** 10.1101/2024.12.18.629266

**Authors:** Paprapach Wongdontree, Milya Palmier, Clara Louche, Vincent Leguillier, Carine Machado Rodrigues, Karine Gloux, David Halpern, Céline Henry, Jamila Anba-Mondoloni, Alexandra Gruss

## Abstract

Phospholipids are essential components of most cell membranes. In *Staphylococcus aureus*, PlsX acyltransferase is considered indispensable for initiating phospholipid synthesis, unless exogenous fatty acids (FAs) are available to bypass this requirement. We report that *S. aureus* can capture internal FA sources to overcome PlsX essentiality in a Δ*plsX* mutant *via* point mutations in either of two genes: *fabF*, which encodes the FA synthesis enzyme 3-oxoacyl-(acyl-carrier-protein) synthase II, or *fadM*, which encodes an understudied bifunctional acyl-CoA thioesterase and ACP binding protein. Despite growth rescue, both Δ*plsX* suppressors differ from the parental strain by producing phospholipids with shortened FA lengths suggesting that both suppressors lead to premature FA release during synthesis. Additionally, both suppressors display increased sensitivity to β-lactam antibiotics. The similar behavior of both suppressors led us to show that *fabF* suppressors require the presence of *fadM*, indicative of FabF-FadM cooperation. We propose that reduced processivity of FabF suppressor variants, or greater availability of FadM for ACP binding in FadM variants, facilitates FA release from FabF-acyl-ACP intermediates. A FabF-FadM relay leading to FA release may contribute to homeostasis between FASII and phospholipid synthesis pathways.

**Significance:** Phospholipids are vital cell membrane components. The essential *Staphylococcus aureus* phospholipid synthesis enzyme PlsX uses acyl-ACP, the end-product of fatty acid (FA) synthesis (FASII), to initiate phospholipid production. Despite its central role, PlsX can be substituted by exogenous FAs whose phosphorylation yields the same product. We discovered that without FA supplementation, mutants arise that rescue growth, indicating that internal FAs are released. Mutations occurred in either FabF, a FASII enzyme, or in FadM, an incompletely characterized protein. Our analyses give evidence that FabF and FadM proteins cooperate, and facilitate FA availability when either protein is mutated. We propose that in normal conditions, FadM might act as an “overflow valve” by releasing FAs from the FabF intermediate, which prevents buildup of FASII intermediates, and ensures FA-phospholipid balance. Remarkably, while this pathway rescues *S. aureus* growth, it sensitizes the MRSA strain to β-lactam antibiotics.

## Introduction

Lipids provide the essential scaffolding for cell membranes and life. Bacterial phospholipid membranes comprise fatty acids (FA) that are attached to a glycerophosphate backbone [1]. The phospholipid FAs are either produced by the FA synthesis (FASII) pathway or captured from the environment [2]. The FASII product, acyl-Acyl Carrier Protein (acyl-ACP or FA-ACP) is a substrate for three enzymes: FabF for FA elongation *via* FASII, and PlsX and PlsC for respectively the first and second steps of phospholipid synthesis (**Fig. 1**).

**Fig. 1:**
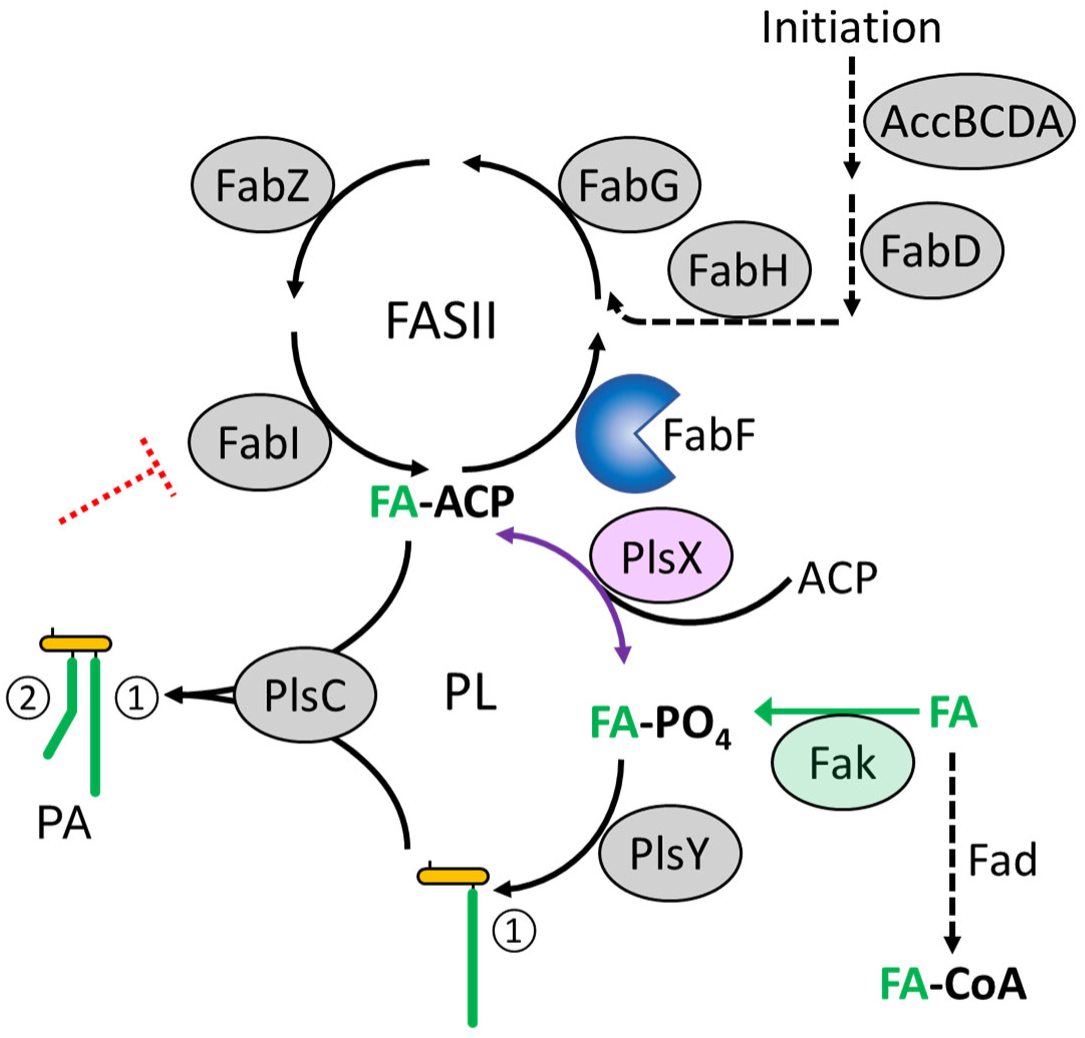
Roles of the reversible PlsX enzyme in phospholipid synthesis. The FASII pathway schematized here produces acyl-ACP (FA-ACP), which may either be elongated by FASII using FabF (blue), or used by PlsX (purple) to initiate phospholipid (PL) synthesis. The PlsX product FA-PO_4_ is a substrate for PlsY, a membrane protein that joins the acyl group to the glycerophosphate backbone in position 1 (①, bottom). PlsC then uses this intermediate product (lysophosphatidic acid), to add the second acyl moiety from FA-ACP to position 2 (②), generating phosphatidic acid (PA, left). Reverse PlsX activity uses phosphorylated exogenous FAs (green) produced by the FA kinase Fak [4], to produce FA-ACP. FASII inhibition (dashed red T) is bypassed by exogenous FAs [6], which are converted to FA-PO_4_. The latter is used by PlsY to fill position 1, and by reverse PlsX, followed by PlsC, to fill position 2, thus producing PA. In a Δ*plsX* mutant, FAs charging position 1 are exogenous, while those charging position 2 are FASII-synthesized; in this case, the two FA sources are disconnected. See [1, 6, 9, 12] for references. Exogenous FA are subject to β-oxidation by FA degradation enzymes particularly in glucose-starved conditions (Fad, lower right; [25, 26, 50]). Enzymes circled in grey are not discussed in this study.

The membrane-associated acyltransferase PlsX catalyzes the first dedicated step of phospholipid synthesis in Bacillota bacteria, by reversibly catalyzing conversion of acyl-ACP, the FASII product, to acyl-phosphate (acyl-PO_4_) [3]. An alternative phospholipid initiation mechanism, not requiring PlsX, involves assimilation of exogenous FAs *via* phosphorylation by the FA kinase Fak [4]. The membrane glycerol-3-phosphate acyltransferase PlsY then transfers the FA moiety of acyl-PO_4_ to the 1-position of the glycerophosphate backbone, forming lysophosphatidic acid. However, reverse PlsX activity may also reconvert acyl-PO_4_ to acyl-ACP when exogenous FAs are available, which then re-enter the FASII system or are used by PlsC for phospholipid synthesis [4, 5] (**Fig. 1**). Providing acyl-ACP by PlsX reverse activity on exogenous FAs, once phosphorylated, allows bacteria to overcome the need for FASII, making them insensitive to FASII inhibitors [6–8]. In *Bacillus subtilis*, PlsX is membrane-associated for phospholipid synthesis, which could facilitate acyl-PO_4_ transfer to PlsY, but likely not for its reverse activity [9]. PlsX is thus a sorting station that initiates phospholipid synthesis, or returns acyl groups for FASII elongation and PlsC processing.

When exogenous FAs are absent, PlsX may still be dispensable in some bacteria. Numerous Pseudomonadota bacteria encode PlsB, which has functional redundancy to PlsX-PlsY [3]. In *E. coli*, suppressors of the synthetic lethal Δ*plsX* Δ*plsY* mutation (despite the existence of the functionally redundant enzyme PlsB [3]) [10] displayed increased pools of glycerol-3-phosphate (Gly-3-P), the backbone for FA attachment to produce phospholipids [10]. Increased Gly-3-P was suggested to compensate the low efficiency acyl-transferase activity of PlsB compared to PlsY. Among Bacillota pathogens, some deploy an acyl-ACP thioesterase (acyl-ACP TE) that liberates endogenous FAs from acyl-ACP, which are then phosphorylated by Fak to initiate phospholipid synthesis [11–14]. Acyl-ACP TE fully restores growth of *Streptococcus pneumoniae* Δ*plsX* [12], and conditionally rescues *Enterococcus faecalis* Δ*plsX* growth; however, the acyl-ACP TE preferentially cleaves acyl-ACP comprising unsaturated FAs, while growth is stimulated by saturated FAs [13]. These alternative systems in Bacillota all generate an internal FA supply to initiate phospholipid synthesis when PlsX function is impaired.

The major human pathogen *Staphylococcus aureus* lacks PlsB, and an acyl-ACP TE to liberate FAs, and only exogenous FAs or a cloned gene encoding heterologous acyl-ACP TE reportedly complemented growth of a *plsX* deletion strain [5]. Our previous work on FASII bypass revealed that exogenous FAs rescue *S. aureus* growth by one of two processes: point mutations that disable *fabD* [7], or by non-mutational adaptation in which FAs are efficiently incorporated after a 6-8 h time lag [6–8]. These studies led us to now investigate the step following FASII, mediated by PlsX. For this we constructed Δ*plsX* mutants, and unexpectedly identified suppressors conferring growth. In this report, we characterized activity-altering mutations mapping to two distinct loci that rescue Δ*plsX* viability without exogenous FAs. Mutations in FASII enzyme FabF, a 3-oxoacyl-ACP synthase II, or in an incompletely characterized acyl-CoA thioesterase and ACP binding protein designated as FadM, rescue Δ*plsX* growth. Our findings reveal that *plsX* essentiality is abolished by point mutations in *fabF* or *fadM*. We demonstrate interdependency of FabF and FadM in rescuing the absence of PlsX. These findings may indicate a general role for FadM as a newly found actor shuttling FAs from FASII towards phospholipid synthesis. Remarkably, the *fabF* and *fadM* suppressors rescue Δ*plsX* growth, but sensitize an MRSA *S. aureus* strain to the β-lactam antibiotic amoxicillin.

## Results

### Construction and confirmation of a Δ*plsX* in-frame mutant

*plsX* is the second gene in a multicistronic operon also comprising FASII genes (**Supplementary Fig. S1**). We constructed in-frame Δ*plsX* mutants of RN-R (RN4220 repaired for a defective *fakB1* gene in the 8325 lineage [15]) and of USA300_FPR3757 JE2 strains. Both Δ*plsX* mutants failed to grow on BHI solid medium in the absence of FAs. Deletions were confirmed by PCR and whole genome sequencing (**Table S1**) and by RN-R Δ*plsX* growth complementation in the absence of FAs using a plasmid-carried copy of the intact *plsX* gene (**Supplementary Fig. S2**).

### FA auxotrophy of *S. aureus* Δ*plsX* mutants is suppressed by a point mutation in *fabF*

During phenotypic characterization of the JE2 Δ*plsX* mutant, we noted the appearance of colonies on BHI medium without added FA (**Fig. 2A**, left plate). Note that BHI reportedly contains traces of FA [16], but which are insufficient to confer Δ*plsX* growth. Two colonies subjected to whole genome sequencing confirmed the in-frame *plsX* deletion and also revealed point mutations mapping to FabF (SAUSA300_0886), the 3-oxoacyl-ACP synthase II of the FASII elongation cycle (see **Fig. 1**), producing FabF^A119E^ (alanine to glutamic acid; Sup1) and FabF^D266A^ (aspartic acid to alanine; Sup2). The suppressor carrying FabF^D266A^ was not recoverable from the primary stocked strain, possibly because the mutation was deleterious for growth. Sup1 and Sup2 also carried the same point mutation in *csa1A* (SAUSA300_0100), encoding a tandem lipoprotein.

**Fig. 2.**
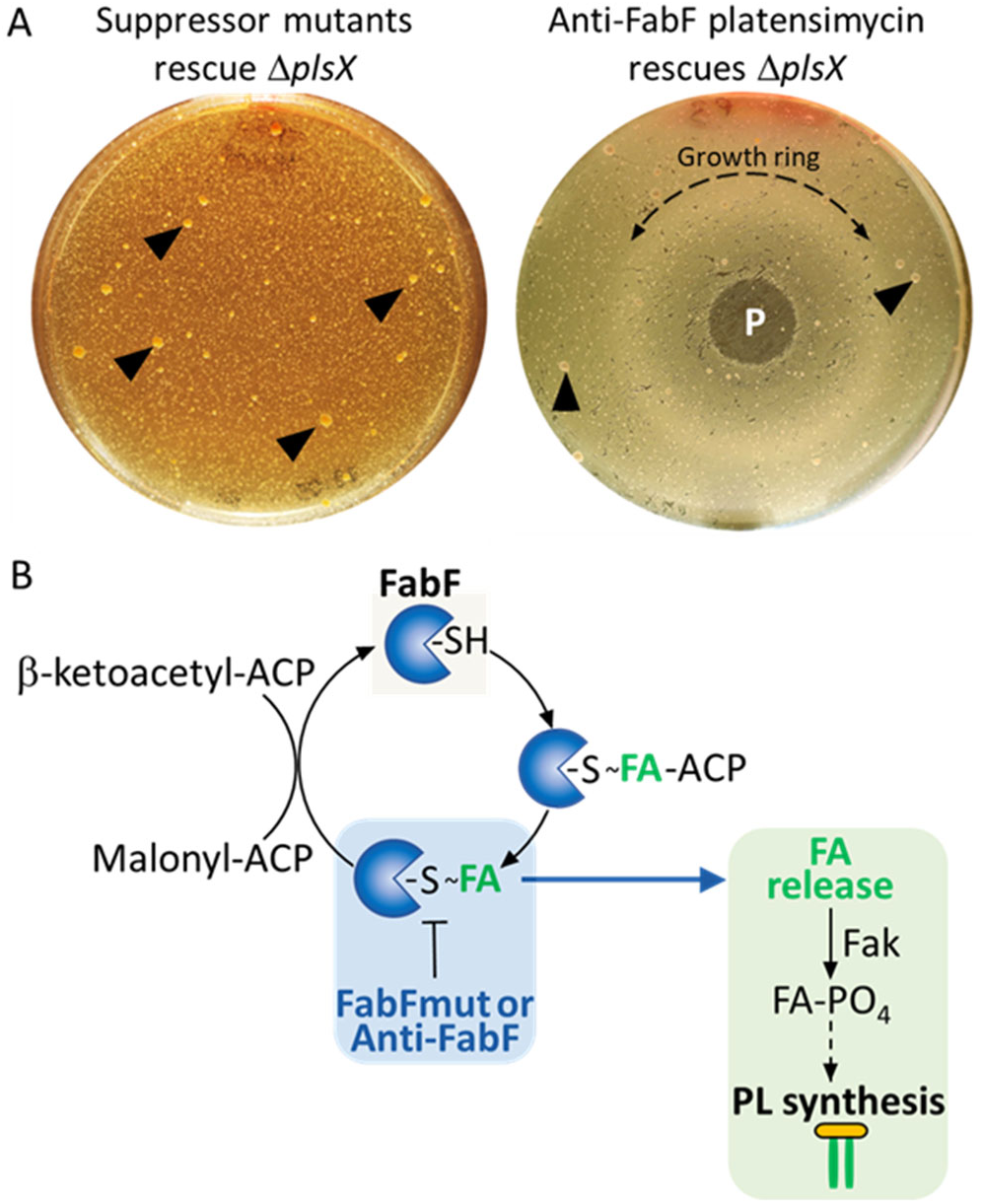
*fabF* mutants or the FabF inhibitor platensimycin rescues Δ*plsX* growth in the absence of exogenous FAs. **A.** Lawns of RN-R Δ*plsX* were spread on BHI medium without FA addition, without (left) or with (right) 1.5 µg platensimycin (P) deposited in the center. Plates were photographed after 4 days at 37°C. Black arrowheads, examples of suppressors, some of which map to *fabF*. Dashed arrow (left), a homogenous growth ring surrounds the platensimycin spot. N=4. **B.** Working model for Δ*plsX* growth rescue by a FabF mutation (FabF mut) or by partial FabF inhibition by platensimycin (anti-FabF). FabF perturbations would reduce enzyme processivity, such that the unstable FabF-FA intermediate releases FAs prior to second substrate malonyl-ACP loading, as needed to bypass the Δ*plsX* defect. FabF in blue; FAs in green. Symbols for FabF are based on [51].

We expanded our analyses by isolating 31 other Δ*plsX* suppressors from RN-R or JE2 backgrounds (16 and 15 suppressors, respectively), and screened for those carrying *fabF* mutations by PCR followed by DNA sequence analyses (**Table S2**). Among them, 14 carried *fabF* mutations, which mapped to the same *fabF* nucleotide position in both strain backgrounds, resulting in a FabF^A119E^ alteration as in Sup1. This allele is referred to as *fabF1*. *csa1A*, which carried a mutation in Sup1 and Sup2 mutants, was not mutated in the 14 *fabF* mutants, indicating that *fabF* was the suppressor allele. Emergence of the same *fabF* mutation in both suppressor strain backgrounds could suggest that few FabF point modifications would allow both FASII activity and Δ*plsX* complementation.

We attempted to prove that the *fabF* mutation was directly responsible for the Δ*plsX* suppressor phenotype by PCR-amplifying the WT *fabF* and mutant *fabF1* alleles, cloning on a medium copy-number vector and constitutive promoter, and establishing plasmids in the Δ*plsX* and/or *fabF1*Δ*plsX* strains, with the goal of reversing the suppressor phenotype. A clone expressing *fabF1*, but not WT *fabF* could be established in the WT RN-R strain by direct transformation.; this is consistent with the putative reduced activity of this allele. We further failed to establish the *fabF1* clone in the Δ*plsX* background, even in FA-supplemented medium. This suggested that *fabF* overexpression was toxic in *S. aureus*, as reported in *Escherichia coli* [17], and that our strategy was not feasible.

As an alternate approach, we used a FabF inhibitor, platensimycin, to determine whether disabling FabF was responsible for Δ*plsX* suppression. This antibiotic binds to the acyl-ACP-FabF intermediate, and competes with malonyl-ACP entry, thus preventing FA elongation [18]. However, it also leads to increased levels of FabF and other FASII enzymes in *Bacillus subtilis* [19]. We tested platensimysin, like the FabF mutant, would overcome the Δ*plsX* growth defect, presumably by slowing FabF activity. RN-R Δ*plsX* cultures were plated on BHI solid medium, and platensimycin (1.5 µg) was deposited on plates, which were then incubated 96 h at 37°C (**Fig. 2A**, right plate). In addition to the emergence of Δ*plsX* suppressors, a distinct homogeneous growth ring appeared around the platensimycin spots. Similar experiments were performed in liquid medium using 2-fold dilutions of platensimycin concentrations ranging from 1 ng to 500 ng/ml (**Supplementary Fig. S3**). Compared to BHI medium, growth between 7 and 12 h was significantly stimulated by the presence of 125 ng/ml platensimycin (P=≤0.05). However, experiments in liquid medium are potentially flawed as mutant emergence and platensimycin stimulation cannot be distinguished. Ring formation on solid medium appears to be a more accurate means of assessing growth stimulation. Since the *fabF* mutants and a FabF inhibitor both restore Δ*plsX* growth, we consider it likely that the FabF mutation is directly responsible for Δ*plsX* suppression. Reducing *fabF* efficacy by mutation or subinhibitory platensimycin treatments could destabilize the FabF-acyl-ACP intermediate, to liberate free FAs and complement the Δ*plsX* growth defect (**Fig. 2B**).

### FA auxotrophy of *S. aureus* Δ*plsX* mutants is suppressed by mutations in *fadM* that map to the acyl-CoA binding cavity

We searched for mutations that conferred Δ*plsX* growth on BHI among the 17 remaining suppressors carrying a wild type *fabF* sequence. Genomic DNA sequencing was performed on three such isolates (one derived from RN-R Δ*plsX* and two from JE2 Δ*plsX* backgrounds). A single common gene target, SAOUHSC_01348 in RN-R or SAUSA300_1247 in JE2, was mutated (**Table S1**, **Table S2**). While annotated as a 1,4-dihydroxy-2-naphthoyl-coenzyme A (DHNA-CoA) thioesterase implicated in menaquinone synthesis, the enzyme was shown to have acyl-CoA thioesterase activity [20]. Moreover, the 155-amino-acid ORF shares ∼40% similarity and a conserved serine, histidine and aspartic acid catalytic triad with FadM, an *E. coli* acyl-CoA thioesterase and ACP binding protein (PBD database protein 1NJK), [20–24]. The *S. aureus* gene, referred to as *fadM*, was PCR-amplified and sequenced in the 17 other suppressors lacking mutations in *fabF*. All carried mutations in *fadM*. A single variant (FadM^Y90F^, [encoded by *fadM2*]) was identified in the four isolates selected from RN-R Δ*plsX*; three distinct FadM variants were identified among the JE2 Δ*plsX* suppressors (FadM^I38T^ [*fadM1*], FadM^Y90F^ [*fadM2*], and FadM^Y133F^ [*fadM3*]) (**Table S1**, **Table S2**). These results point to FabF and FadM as the main suppressors rescuing Δ*plsX* growth.

The *S. aureus* FadM crystal structure (PDB: 6FDG) and Alphafold predictions (https://alphafold.ebi.ac.uk/entry/A0A0H2XFE3) provide detailed information for mapping the identified FadM variants conferring Δ*plsX* suppression. The variants coincided with or mapped adjacent to amino acids reportedly involved in forming an FA binding cavity that favors long FA substrates [20] (**Fig. 3A**). The grouped location of these variants leads us to speculate that acyl-CoA thioesterase activity may be defective in the *fadM* suppressors.

**Fig. 3.**
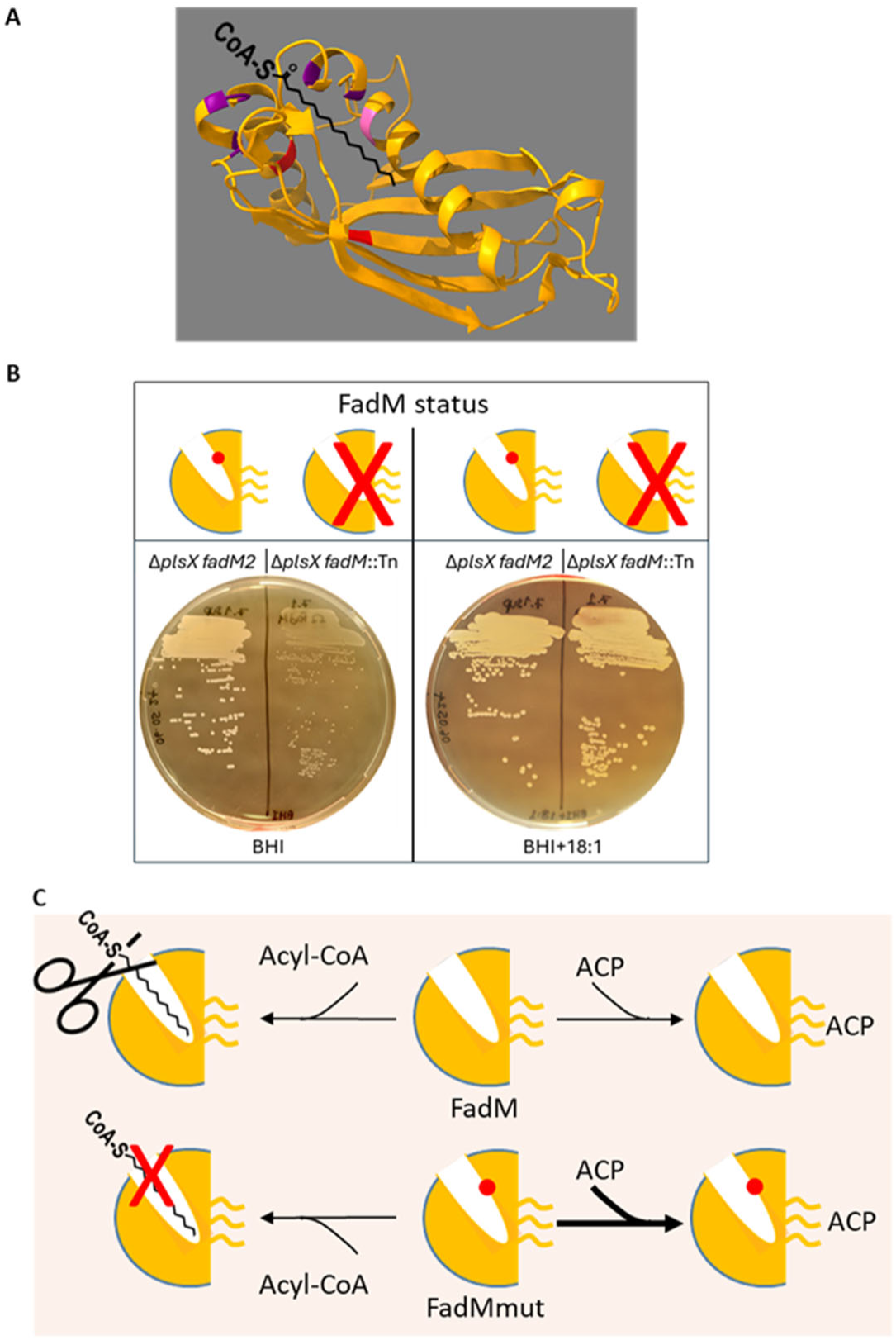
FadM variants mapping to the predicted FA binding tunnel, but not FadM inactivation, confer the *fadM* suppressor phenotype. **A.** Schematized *S. aureus* FadM monomer binding to FA, based on crystal structure and Alphafold predictions ([20, 37], designed with https://alphafoldserver.com and http://www.cgl.ucsf.edu/chimera [52]). FA moiety (black line) binding in the FadM cavity involves amino acids Ile38 (pink), Tyr45; Met48, Leu122, Tyr125, and Phe126 (purple) [20]. The Δ*plsX* suppressors mapping to FadM affect amino acids Ile38Thr, Tyr90Phe, or Tyr133Phe (pink, and 2 red, respectively), which cluster around the FA-binding cavity; Ile38 (pink) is common to structural predictions and the FadM^I38T^ Δ*plsX* suppressor. **B.** FadM inactivation abolishes Δ*plsX* suppression: Growth of Δ*plsX fadM2* and the Δ*plsX fadM::*Tn derivative was compared on solid BHI medium without or with C18:1. Plates were photographed after 48 h growth at 37°C. Background growth on BHI (left) may be due to FA carryover from pre-cultures or to FA traces in medium. Plates are representative of biological triplicates. **C.** Working model for FadM *versus* FadM mutant function. Binding of the acyl-CoA FA moiety (zigzag line) in the FadM tunnel is needed for acyl-CoA thioesterase activity (scissor). FadM mutations in the tunnel (red dot) prevent this activity (red X) but conserve ACP binding (thick arrow, lower right).

If FadM acyl-CoA thioesterase activity were needed for Δ*plsX* suppression, then its substrate, acyl-CoA, would need to be produced. In *S. aureus*, acyl-CoA is synthesized by acyl-CoA synthetase (encoded by *fadE*), which is part of the recently elucidated FA degradation (*fad*) system (**Fig. 1**; [25, 26]). The *fad* genes are subject to CcpA-mediated glucose repression [25–27]. However, glucose did not affect the *fadM* mutant Δ*plsX* suppressor phenotypes (**Supplementary Fig. S4**). Based on above results, we consider it reasonable to rule out a role for FadM acyl-CoA thioesterase activity in Δ*plsX* growth rescue.

### A *fadM* null mutant does not confer Δ*plsX* growth

Suppression by *fadM* point mutants could be due to FadM-mediated ACP binding, or to the loss of all FadM activity. We asked whether inactivation of *fadM*, the downstream gene in a 2-gene operon comprising *acnA* ([27]; **Supplementary Fig. S1**), suppresses the Δ*plsX* phenotype. For this, a *fadM* transposon insertion from the Nebraska Transposon Mutant Library library, interrupting FadM at amino acid position 5, was transferred by ϕ80-mediated transduction into the JE2 Δ*plsX fadM2* suppressor (strain 7.1; **Table S2**) [28]. The Δ*plsX fadM::*Tn strain failed to grow on FA-free medium (**Fig. 3B**). Since total loss of FadM does not promote Δ*plsX* suppression, we conclude that some FadM activity is required for *S. aureus* Δ*plsX* suppression. As FadM I38T, Y90F, and Y133F mutations each map to the acyl-CoA thioesterase binding tunnel, we propose that the FadM ACP binding activity is involved in Δ*plsX* suppression (**Fig. 3C**).

### Impact of Δ*plsX* suppressors on bacterial growth properties

Growth of Δ*plsX* suppressors was assessed in RN-R and JE2 backgrounds in non-FA-supplemented BHI medium (**Fig. 4A**). The initial Δ*plsX* deletion strains fail to grow unless supplemented with FAs. Both *fabF* and *fadM* suppressor mutants grew similarly, but their growth phenotypes differed according to the parental strains: in the RN-R background, growth of both tested suppressors was intermediate between the WT strain and the Δ*plsX* mutant. In contrast, JE2 derivative suppressors grew like the parental strain. A distinguishing feature between the two *S. aureus* strains is the production of staphyloxanthin pigment by JE2 but not by RN-R (an RN4220 derivative), whose hydrophobic long-chain isoprenyl moiety of staphyloxanthin [29] could conceivably stabilize the membrane in suppressor strains. However, reduced pigmentation of Δ*plsX*, and both *fabF* and *fadM* suppressors compared to WT and *fadM::*Tn controls (**Fig. 4B**), indicate that other factors are involved in improved rescue in the JE2 background.

**Fig. 4.**
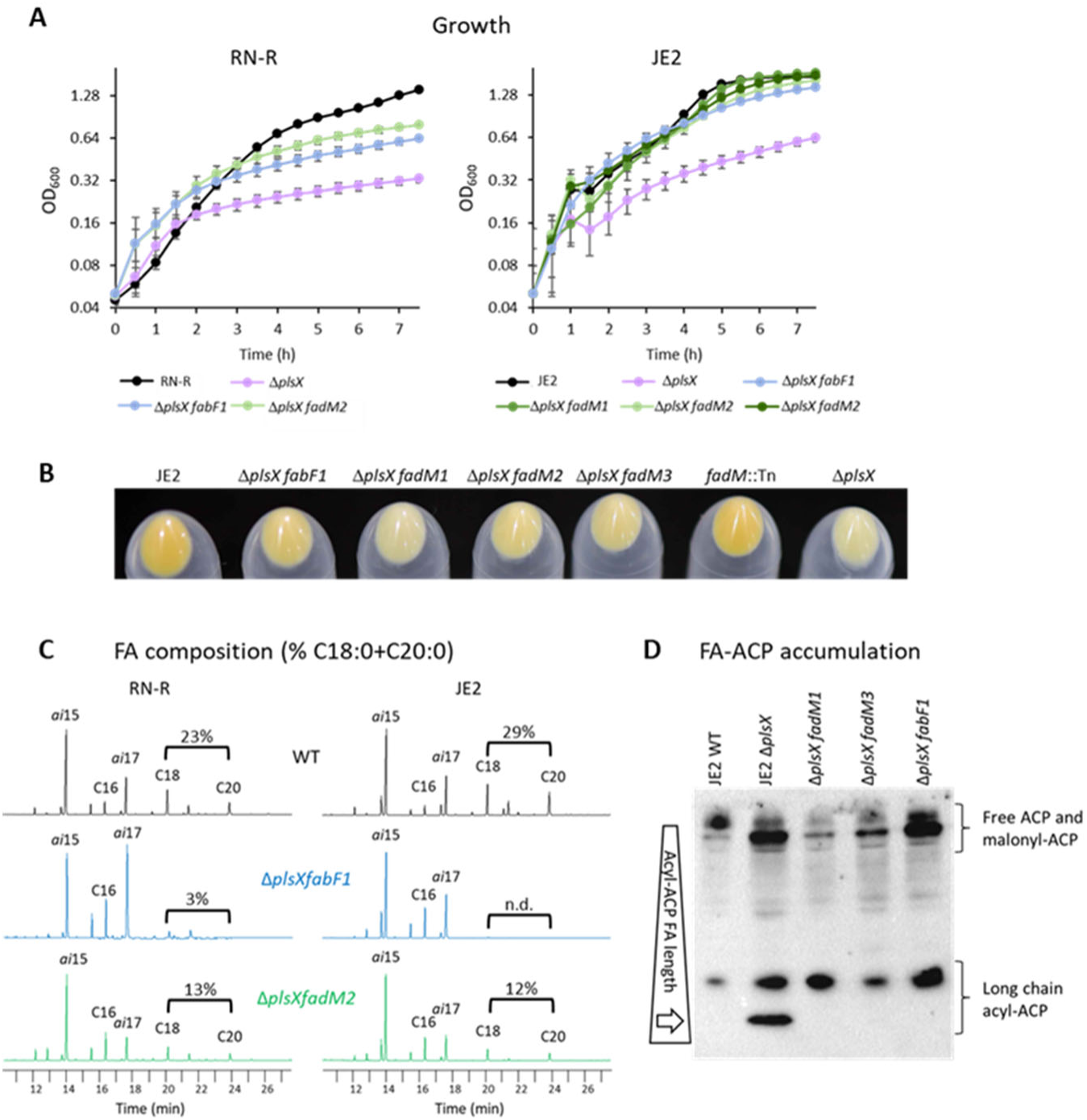
Behavior of *fabF* and *fadM* suppressors of Δ*plsX* without exogenous FAs. **A.** Growth of WT *S. aureus* (RN-R or JE2), Δ*plsX*, and Δ*plsX fabF* or Δ*plsX fadM* suppressors in BHI medium. Left: RN-R WT, and derivatives: Δ*plsX*, and Δ*plsX* suppressors *fabF1* (encoding FabF^A119E^) and *fadM2* (encoding FadM^Y90F^). Right: JE2 WT, and derivatives: Δ*plsX*, and Δ*plsX* suppressors *fabF1*, *fadM1*, *fadM2*, and *fadM3* (encoding FabF^A119E^, FadM^I38T^, FadM^Y90F^, and FadM^Y133F^ respectively). Mean and standard deviation of independent triplicate cultures are shown. **B.** BHI + C18:1-grown overnight cultures of the indicated strains were pelleted, and photographed to compare pigmentation. C18:1 was added to include Δ*plsX* in comparisons. **C.** FA profiles of RN-R (left)- and JE2 (right)-derived WT, Δ*plsX*, and *fabF* or *fadM* suppressors determined from BHI-grown cultures. FA assignments are as indicated. *ai*, anteiso; even numbered FAs are saturated (e.g., C16 is C16:0). The main FA component of WT *S. aureus* is *ai*15 [7]. Peak heights correspond to relative responses (mV) of each FA in a sample. The proportions (in %) of C18:0 plus C20:0 of total FAs are indicated over the brackets. **Table S3** provides complete FA profile information. **D**. Acyl-ACP species in *S. aureus* JE2-derived Δ*plsX*, and Δ*plsX* suppressor strains. JE2 WT, Δ*plsX fadM1*, Δ*plsX fadM3*, and Δ*plsX fabF1* (isolates 7.5, 8.7, and 7.7, respectively; **Table S2**) were grown in BHI. The Δ*plsX* strain was pre-grown in BHI+C18:1 medium, and FA-starved for 2h in BHI prior to harvesting. Extracts were loaded on non-denaturing acrylamide gels, and immunoblotted using anti-ACP [7, 35, 53]. Longer FA moieties on acyl-ACP migrate faster on gels (wedge; [35, 54]). White arrow, accumulated acyl-ACP produced by FASII is not processed to phospholipids [5] (Fig. 1). Results are representative of biological duplicates.

### Membrane FAs are shortened in *fabF* and *fadM* suppressors of Δ*plsX*

We considered that the *fabF1* point mutation in the Δ*plsX fabF1* suppressor strain might alter FabF processivity, suggesting a shift in FASII products. We also asked whether the mutations in *fadM,* encoding a possible FabF partner, might have similar effects. We therefore assessed membrane FA composition of both Δ*plsX* suppressors in RN-R and JE2 backgrounds in medium devoid of FAs (**Fig. 4C**, **Table S3**). The *fabF* suppressors produced membrane FAs of markedly shorter overall length in both strain backgrounds. Notably, straight-chain FAs C18:0 and C20:0, represent 23-29% of total FAs in parental strains but are essentially absent in *fabF* suppressors. This suggests that processivity of FabF^A119E^ expressed from the *fabF1* suppressor allele is reduced compared to WT FabF. In *S. aureus*, branched chain precursors are preferred substrates for FabH (see **Fig. 1**; [30]), which might lead to higher proportions of branched chain FAs (*ai*15 and *ai*17) in *fabF1* suppressors. FA profiles in the *fadM2* suppressor showed a 2-fold decrease in proportions of C18:0 and C20:0 compared to those in the respective parental strains (**Fig. 4C**, **Table S3**). These results confirm that both *fabF* and *fadM* mutant suppressors of Δ*plsX* lead to decreased proportions of long chain saturated FAs in phospholipids.

### Δ*plsX* suppressors display accrued β-lactam sensitivity

β-lactam resistance of methicillin resistant *S. aureus* (MRSA) is associated with the presence of penicillin binding protein Pbp2A’ (encoded by *mecA*), and is a main cause of treatment failure [31, 32]. Interestingly, disturbance of membrane lipid microdomains affects Pbp2A’ oligomerization and compromises resistance [33]. We asked whether the membrane changes induced by Δ*plsX* suppressors, notably their shorter FAs (**Fig. 4C**) might also impact β-lactam antibiotic resistance profiles. For this, amoxicillin minimal inhibitory concentrations (MICs) were measured by Etests. Tests were performed on BHI solid medium without and with 2.5 µM C18:1; the FA addition allowed us to include Δ*plsX* in these comparisons (**Supplementary Fig. S5**). Control strains (WT, *fadM::*Tn, and Δ*plsX*) showed comparable resistance to amoxicillin in both media (0.25-0.38 µg/ml by eTest). Remarkably, all Δ*plsX* suppressors, carrying *fabF* or *fadM* mutant alleles, showed accrued sensitivity, with similar antibiotic resistance phenotypes (0.064-0.094 µg/ml).

### Acyl-ACP species vary in JE2 WT, Δ*plsX*, and Δ*plsX* suppressors

Since PlsX uses acyl-ACP, the FASII end product, to initiate phospholipid production, its deletion provokes acyl-ACP accumulation (**Fig. 1**; [5, 34]). Exogenous FA addition resolves accumulated acyl-ACP in a Δ*plsX* mutant [12]. We assessed acyl-ACP intermediate accumulation in the JE2 Δ*plsX* mutant and derivative suppressors by non-denaturing polyacrylamide gel electrophoresis, and probing with anti-ACP antibodies [7, 35] (**Fig. 4D**). Unlike the WT strain, the Δ*plsX* mutant was distinguished by the presence of a fast-migrating band corresponding to ACP bound to long-chain FA generated by FASII, likely the dominant FA *ai*15. The Δ*plsX* strain assimilates acyl-ACP to form phosphatidic acid (PA) only if an exogenous FA (e.g. C18:1) is added to complete PA synthesis (**Fig. 1**). The long acyl-ACP species was absent in Δ*plsX fabF1* and Δ*plsX fadM1, M2*, or *M3* suppressors. This observation, and good growth of suppressor strains (**Fig. 4A**) indicate that acyl-ACP is processed to initiate phospholipid synthesis and bypass the phospholipid synthesis block in Δ*plsX*.

### Δ*plsX* suppressors do not compensate reverse PlsX activity, which is required to bypass the FASII pathway

The above results show that *fabF* and/or *fadM* suppressors compensate the absence of PlsX in normal growth conditions, i.e., in converting acyl-ACP into acyl-PO_4_. We asked whether *fabF* and/or *fadM* suppressor strains could also complement the reverse PlsX activity, i.e., conversion of acyl-PO_4_ to acyl-ACP (**Fig. 1**, **Fig. 5A**). If this were the case, FASII inhibition in Δ*plsX* suppressors would be bypassed by FA supplementation to growth medium. To test this, WT strain JE2, and isogenic Δ*plsX*, Δ*plsX fabF1*, and Δ*plsX fadM2* strains were grown in FA-supplemented medium (SerFA, containing 250µM of an FA mixture and 10% fetal calf serum; see Materials and Methods), without and with the FASII inhibitor AFN-1252 [36] (**Fig. 5B**). All strains grew to equivalent densities in the absence of inhibitor. The JE2 WT strain overcame the presence of AFN-1252 in SerFA [6, 8]. In contrast, neither the Δ*plsX* mutant nor the Δ*plsX* suppressors could bypass the FASII inhibitor. These findings show that *fabF* or *fadM* suppressor mutations do not compensate the PlsX reverse reaction, and that FASII must be active for Δ*plsX* suppressor growth.

**Fig. 5.**
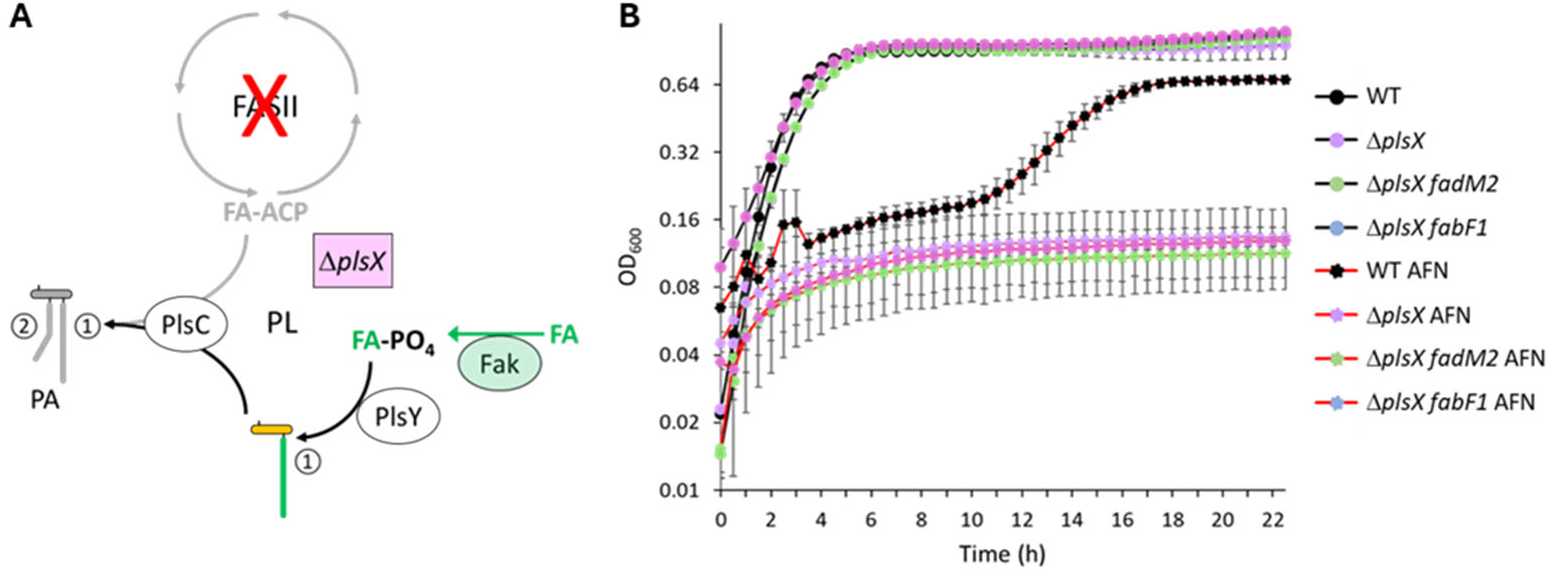
Synthetic lethal effect of a combined FASII and PlsX block in Δ*plsX* and suppressor strains. **A.** Expected effects of FASII inhibition on an *S. aureus* Δ*plsX* mutant. In a WT strain, FASII inhibition by mutation or by antibiotic action is overcome by exogenous FAs, which are phosphorylated by Fak [4]. FA-PO_4_ is the substrate for PlsY and for reverse PlsX activities, and is thus used for phospholipid (PL) synthesis at both the 1 and 2 positions. If both FASII and PlsX are blocked as shown, exogenous FAs can only fill position 1. The FASII defect or PlsX absence can be rescued separately by FA supplementation, but not the combination. Green, exogenous FAs and FA kinase; purple, *plsX* deletion; grey, pathways and products affected by combined FASII and PlsX block. **B.** Effects of anti-FASII treatment on JE2 Δ*plsX*, and Δ*plsX* suppressor strains in FA-supplemented medium. Strains were precultured in SerFA, and grown in SerFA without or with addition of 0.5 µg/ml anti-FASII AFN-1252. When the anti-FASII is added, WT *S. aureus* strains incorporate FAs to constitute membrane phospholipids, thus bypassing inhibition [6, 8, 55]. As Δ*plsX* suppressors cannot generate FA-ACP needed for FASII bypass [6, 9, 12], they remain sensitive to anti-FASII. Mean and standard deviation of independent triplicate cultures are shown. Black lines, no anti-FASII, red lines with anti-FASII. *fadM2* encodes FadM^Y90F^; *fabF1* encodes FabF^A119E^.

As the Δ*plsX fabF* and *fadM* suppressors behaved similarly in all the above phenotypic tests (Figs. 4,5, S4, and S5), we considered that these alleles might cooperate in a single pathway.

### FadM is required for emergence of Δ*plsX* suppressors mapping to FabF

As *fadM* interruption abolished Δ*plsX* suppression (**Fig. 3**), we reasoned that suppressors arising in the Δ*plsX fadM::*Tn mutant would map to *fabF*. This hypothesis was tested by comparing suppressor emergence in JE2 Δ*plsX* and Δ*plsX fadM::*Tn backgrounds on solid medium without FA supplementation. We also tested for a stimulatory effect of the FabF inhibitor platensimycin as seen for the Δ*plsX* mutant (**Fig. 2**). Unexpectedly, no suppressors arose after extended (> 1 week) plate incubation at 37°C, compared to frequent mutant emergence in the Δ*plsX* background after 1-2 days (**Fig. 6A, upper**). Moreover, no stimulatory growth ring was detected surrounding the platensimycin spot (**Fig. 6A, lower**). These results affirm that FadM is needed to obtain FabF suppressors of Δ*plsX*. Finally, we introduced the *fadM::*Tn allele into the Δ*plsX fabF1* suppressor mutant. Compared to good growth of Δ*plsX fabF1*, introduction of the *fadM::*Tn allele arrested grow in the absence of exogenous FA, proving that *fadM* is required for *fabF1* rescue (**Fig. 6B**).

**Fig. 6.**
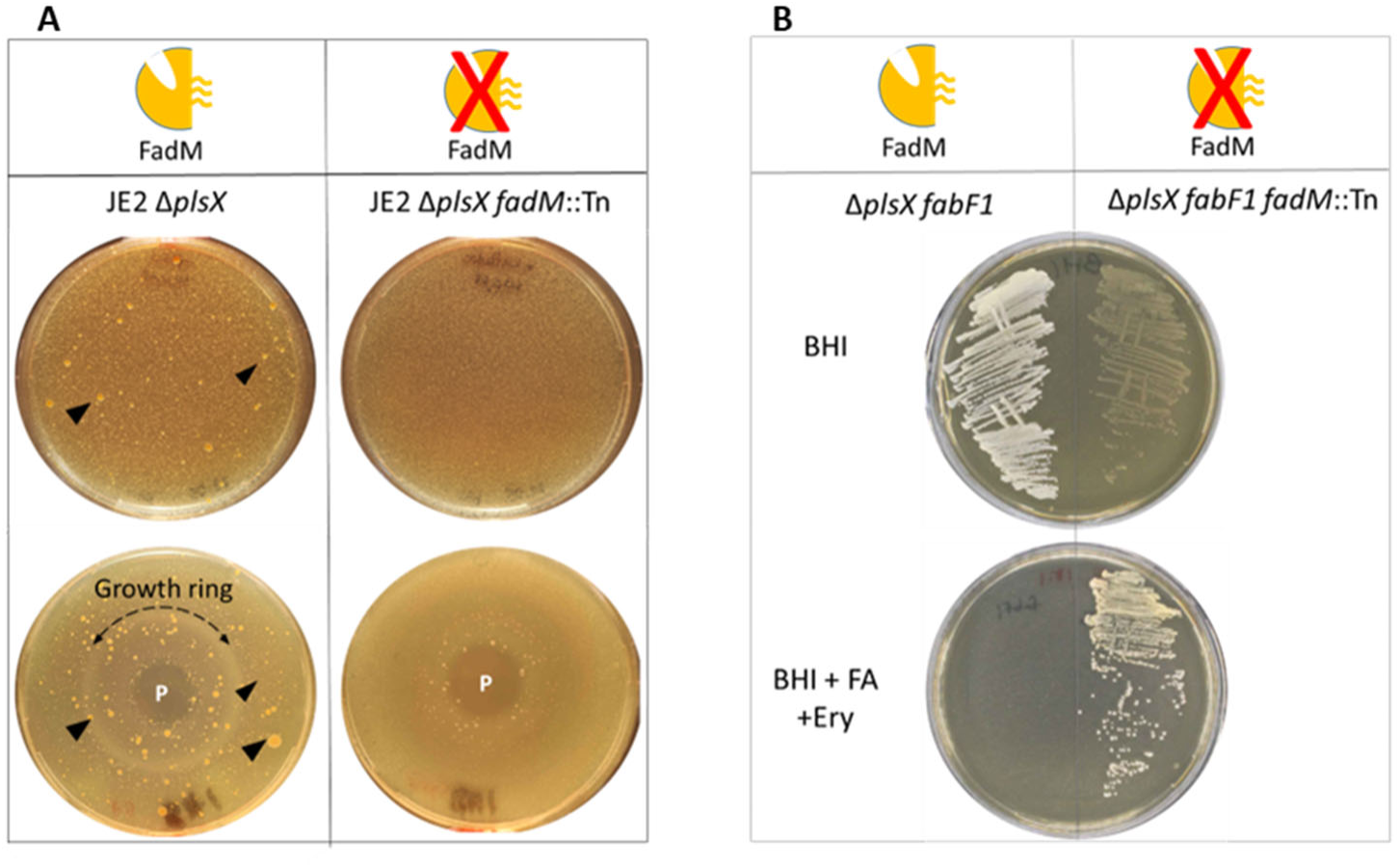
*fabF* suppressors of Δ*plsX* require the presence of a *fadM* allele. **A.** FadM is present in JE2 Δ*plsX* (left) and absent in Δ*plsX fadM::*Tn (right, constructed from strain 7.1: **Table S2**). Strains were plated on BHI solid medium (100 µl of OD_600_ = 0.005), and in the lower set, spotted with platensimycin (P; 1.5 µg). Plates were photographed after 6 days at 37°C. Black arrowheads point to Δ*plsX* suppressor clones. The dashed curved line indicates the growth ring surrounding platensimycin spotting. The small colonies surrounding platensimycin on Δ*plsX fadM::*Tn lawns (right) failed to regrow on BHI medium. **B.** FadM is present in the JE2 Δ*plsX fabF1* suppressor strain (left) and absent in Δ*plsX fabF1 fadM::*Tn (right, constructed from strain 7.8: **Table S2**). Strains were streaked on BHI solid medium (upper), or BHI medium containing 250 µM C18:1 and 5 µg/ml erythromycin (Ery; lower). Plates were photographed after 24 h incubation at 37°C. Results are representative of three independent cultures and platings.

The potential relevance of FabF-FadM cooperation was examined in a WT background by comparing platensimycin inhibition zones in the WT and *fadM::*Tn derivative on BHI medium (**Supplementary Fig. S6**). While inhibitory zones were equivalent, the WT strain displayed a distinct growth ring (24 h incubation) that is not formed in the *fadM::*Tn mutant. Growth ring rescue from the anti-FabF platensimycin inhibitor also requires FadM in the WT background. Based on all the above findings, we conclude that FadM and FabF have cooperative roles.

## Discussion

*S. aureus* requires the phospholipid synthesis enzyme PlsX for growth when FAs are not available in its environment. Here, we show that mutations affecting either of two proteins, FabF (a FASII core enzyme), or FadM (a bifunctional protein with acyl-CoA thioesterase and ACP binding activities), allow *S. aureus* to grow even without this essential enzyme [20, 23, 27]. To explain growth rescue, we propose that the FabF^A119E^ variant Is responsible for premature release of the FA-ACP intermediate. This explanation is consistent with the shortened FAs in membrane phospholipids and with Δ*plsX* growth stimulation by low anti-FabF concentrations. The FadM I38T, Y90F, Y133F variants all map to the acyl binding tunnel as based on crystallographic and biochemical evidence, and likely impair acyl-CoA thioesterase activity [20, 37]. A functional connection is established between FabF and FadM, as both alleles must be present in order to obtain Δ*plsX* suppressors. Interestingly, all tested phenotypes of the *fabF* and *fadM* suppressors, e.g., growth kinetics, membrane FA shortening, and antibiotic sensitivities, were similar. While the nature of FabF-FadM cooperation, whether direct or indirect, remains to be elucidated, these results identify FadM as a key modulator linking FASII and phospholipid homeostasis in *S. aureus*.

We propose a working model to explain Δ*plsX* growth rescue by FabF and FadM variants [38] (**Fig. 7**). In this model, the FabF^A119E^ variant (or FabF in the presence of subinhibitory amounts of platensimycin) may stall and release its FA-ACP cargo before malonyl-ACP entry, which is needed for FA elongation. ACP binding by FadM would facilitate ACP removal from the FabF cavity, thus favoring FA release. The FA made available would then be phosphorylated by Fak for entry into the phospholipid synthesis pathway.

**Fig. 7.**
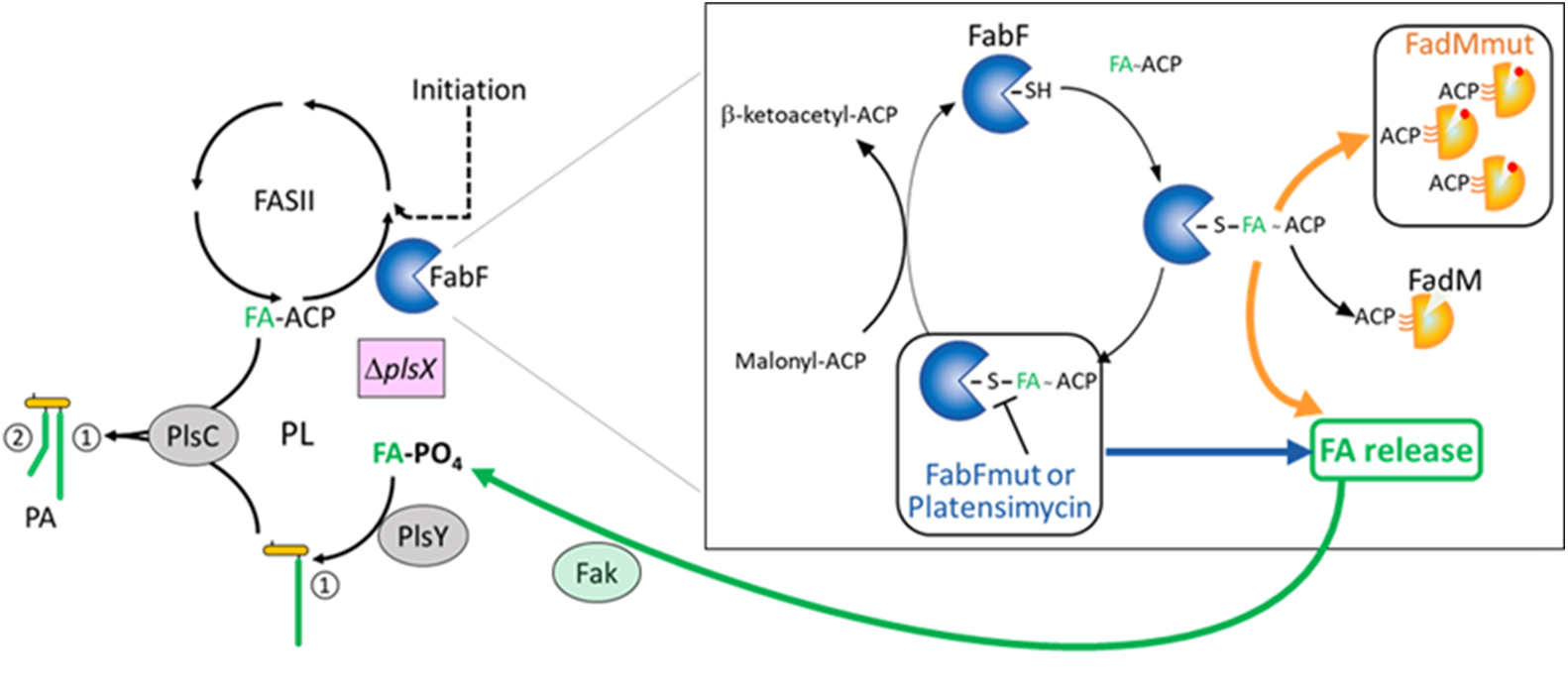
Proposed role of FabF - FadM interactions in controlling FA release and recycling from FabF intermediates. Our study showed epistasis between FabF and FadM, and leads us to a working model of how these proteins interact (depicted in the inset). Left, FASII and phospholipid synthesis (PL) coupled pathways [1, 56]. The *plsX* deletion dissociates the pathways, such that exogenous FA is required for survival. Symbols are as in Fig. 1. **Inset:** FabF - FadM interactions coordinately overcome Δ*plsX* growth arrest when either function is mutated. In the first FASII step, FabF (blue packman and blue arrow) cleaves acyl-ACP (FA-ACP) to release ACP, which then allows malonyl-ACP entry into the FabF cavity to extend the acyl intermediate by 2 carbons [18, 38]. The FabF^A119E^ variant (FabFmut), or addition of anti-FabF inhibitor platensimycin, impedes malonyl-CoA entry, leading to FA release (FA symbols in green). FadM (gold) is bifunctional, with acyl-CoA thioesterase and ACP binding activities [20, 22–24]. Suppressor mutations in FadM (FadMmut red dot) alter the FadM acyl binding cavity, which we propose decreases acyl-CoA thioesterase activity as reported for FadM mutants also mapping to the binding cavity [20]. More FadMmut is thus available for ACP binding (boxed FadMmut). FadM-ACP or FadMmut-ACP binding could favor ACP release from the FabF pocket and leakage of FAs, which are then processed for PL synthesis *via* FA kinase (Fak) [4]. Symbols for FabF are based on [51].

FadM suppressor variants with modifications in their FA-binding cavity are predicted to exhibit reduced acyl-CoA thioesterase activity, as previously observed [20] and proposed here. This would increase the availability of FadM to facilitate ACP removal and FA release from the FabF intermediate. In contrast, a FadM null mutation would exclude this process, consistent with our findings that *fadM* is required for Δ*plsX* suppression.

The newly uncovered contribution of FadM to FabF activity may have relevance to WT *S. aureus* FASII-phospholipid homeostasis by acting as an FA release valve that decreases FabF processivity, leading to shortened phospholipid FAs, e.g., under stress conditions. This hypothesis is currently being investigated.

Remarkably, the Δ*plsX* suppressor strains are β-lactam-sensitive as compared to the MRSA JE2 parent or its Δ*plsX*, and *fadM::*Tn derivatives (**Supplementary Fig. S5**). Both *fabF* and *fadM* suppressors display similar greater amoxicillin sensitivity, as expected if FabF and FadM act on the same pathway. The mechanisms underlying β-lactam-sensitivity and the potential role of membrane synthesis or composition remain unclear, but may involve destabilization of PBP2’, as reported [33]. Thus, rerouting phospholipid synthesis rescues *S. aureus* Δ*plsX* growth but reduces the bacterial response to β-lactam antibiotics. These observations open perspectives for understanding how membrane perturbations, as occur in Δ*plsX* suppressor strains, might resensitize bacteria to this important class of antibiotics.

## Materials and methods

### Strains, growth conditions, and plasmids

Strains and plasmids are listed in **Table S4**. *S. aureus* strains are derived from USA300_FPR3757 strain JE2 [28] and RN4220-R (called here RN-R; [15]). RN-R is a readily transformable strain derived from RN4220, of the NCTC 8325 lineage [39]. It was repaired to restore a functional *fakB1* gene, which is defective in the entire NCTC 8325 lineage [15]). *S. aureus* parental strains and Δ*plsX*-suppressor mutants were grown in BHI medium (Brain Heart Infusion; Gibco, France) or in LB (Lysogeny broth; [40]) as specified, at 37°C with shaking. Medium was supplemented with C18:1 (oleic acid) at a concentration of 250 µM (Larodan, Sweden) when indicated. The FabI inhibitor AFN-1252 (Med ChemExpress, France) and FabF inhibitor platensimycin (Bioaustralis-Tebubio France) were used as specified [18, 41].

### Δ*plsX* mutant construction

The Δ*plsX* mutants were constructed in RN-R (RN4220 repaired for a defective *fakB1* gene in the NCTC8325 lineage; [15]) and JE2 backgrounds. The preparatory plasmid for *plsX* deletion was constructed using the Gibson assembly method to insert DNA segments flanking the *plsX orf* into Sma1-linearized pMAD, a thermosensitive plasmid expressing erythromycin resistance [42, 43]. Oligonucleotides for Δ*plsX* construction are in **Table S5**. Constructions were verified by DNA sequencing of PCR-amplified fragments. The double cross-over gene replacements were obtained as described [15, 43] in RN-R and JE2 strains. Deletions removed nucleotides 1,228,616-1,229,594 in JE2, and 1,148,670-1,149,648 in RN-R, as per respective annotated genomes. The resultant strain genotypes were confirmed by whole genome sequencing (Eurofins, Germany).

### *plsX* cloning and complementation of the Δ*plsX* mutant in RN-R and JE2 strains

To confirm that strain characteristics were due to the *plsX* deletion, the intact *plsX* gene was cloned with its natural promoter (upstream of *fapR*) on multicopy plasmid pIM-locus1 (a pIMAY derivative [44, 45]). Cloning was performed by Gibson assembly with oligonucleotide primers (**Table S5**).

### Isolation of suppressor mutants

RN-R Δ*plsX* and JE2 Δ*plsX* strains were first streaked on solid BHI medium containing 250 µM C18:1, an FA that promotes growth. For each strain, a single colony was used to inoculate a liquid culture (in BHI+C18:1). When the culture reached exponential phase, bacteria were centrifuged, resuspended in BHI, and adjusted to OD_600_ = 0.1. Dilutions were plated on solid medium to obtain ∼10^5^ to ∼10^7^ colony forming units per plate on solid BHI medium without FAs. Colonies were scored after two or more days of incubation at 37°C, and selected at random for restreaking on solid BHI medium. Liquid cultures were prepared in BHI, and stocked in 20% glycerol for storage at -70°C. Selected mutant genotypes were confirmed by whole genome sequencing or by PCR fragment sequencing (Eurofins, Germany) to identify mutated loci.

### Genomic DNA preparation for whole genome sequencing

Bacterial cell pellets, prepared from 3 mL liquid exponential phase cultures, were centrifuged at 8,000 rpm for 5 min. Pellets were re-suspended in 800 µL Tris-EDTA buffer (10 mM Tris-HC1, pH 8.0/1 mM EDTA) plus 20 µL lysostaphin 5 mg/mL, and incubated for 45 min at 37°C. DNA was prepared by DNeasy Blood & Tissue Kit (Qiagen, France) and samples were outsourced for whole genome DNA sequencing using 2x150 bp paired end chemistry and bioinformatics analyses (Eurofins Genomics, France) [6]. SNPs that differed in non-antibiotic-treated JE2 and RN-R strains from those in the reference sequence (GenBank Nucleotide accession codes NC_007793.1 and NC_007795.1 respectively) were subtracted prior to variant identification. Variants were identified as representing ≥80% of reads in sequences for which there were at least 100 reads.

### PCR amplification and confirmation by DNA sequencing

Selected candidate colonies were suspended in 100 μL of 0.9% NaCl with 1 μL of lysostaphin from a 5 mg/mL stock. Tubes containing suspensions were shaken for 45 min at 37°C. Contents were then transferred to tubes containing glass beads and samples were vortexed for 1 min. After a 30 second centrifugation at 8000 rpm, PCR amplification was performed on 1 µl of supernatant, using primers (**Table S5**) and Phusion DNA polymerase (ThermoFisher Scientific, France) according to manufacturer protocol, using a SimpliAmp Thermal Cycler (ThermoFisher Scientific, France). PCR products were analyzed on a 0.8% agarose gel prepared in 40 mM Tris acetate 1 mM EDTA buffer containing 0.5 μg/mL ethidium bromide. Gels were run at 100 V for 30 min in an electrophoresis chamber (Mupid ONE; Dutscher, France), and photographed under UV light using the EBOX VX5 imaging system (Vilber Lourmat, France). PCR samples were purified using the NucleoSpin® Gel and PCR Clean-up kit (Macherey-Nagel, France), and sent for DNA sequencing (Eurofins, France).

### Effects of the FabF inhibitor platensimycin on Δ*plsX* and Δ*plsX fadM::*Tn strains

Strains were grown in BHI containing C18:1 (250 µM) overnight, washed once in BHI medium, and resuspended in BHI to a final density of OD_600_ = 0.1. From this, 100 µl of the bacterial suspension was plated on solid BHI medium. Platensimycin (1.5 µg in 3 µl) was spotted in the center of plates, which were then incubated at 37°C for 4-7 days and photographed.

### Effect of glucose on growth of Δ*plsX* suppressors

Strains were precultured in liquid LB (n.b., LB lacks glucose) or LB with C18:1 for Δ*plsX* at 37°C. After overnight growth, cultures were diluted to OD_600_ = 0.05 and grown for 3-4 hours. Δ*plsX* cultures were first centrifuged and resuspended in fresh LB medium to remove free FAs prior to regrowth. Bacterial densities were then adjusted to OD_600_ = 1. Cultures were streaked on solid medium, corresponding to LB or LB containing 0.5% glucose, which represses *fad* operon expression [25, 27]. Plates were incubated at 37°C for 24 hours and photographed.

### Growth kinetics

Three mL precultures were prepared in BHI for parental and Δ*plsX* suppressor strains, or in BHI containing 250 µM C18:1 for Δ*plsX* mutants. After overnight growth, culture OD_600_ were adjusted to OD_600_ = 0.05 in 200 µL BHI medium without or with C18:1 (250 µM), in 96-well microtiter plates. Growth was followed on a Spark M10 plate reader (Tecan, France). For experiments in which samples were taken for FA determinations or for Western blot experiments, cultures were prepared in 3 to 10 ml liquid cultures, and were monitored manually by OD_600_ measurements every two hours. Amoxicillin sensitivity was determined using AMOXICILLIN AC 256 eTests (BioMérieux, France).

### Determination of membrane FA profiles

Bacterial cultures (equivalent of OD_600_ = 2) were centrifuged, and pellets were washed once in 1 ml of 0.02% Triton in 0.9% NaCl, then twice with 1 ml of 0.9% NaCl. FA extractions were performed as described [46] [6]. Briefly, 0.5 mL of 1N sodium methoxide in methanol was added to bacterial pellets in 2 ml microfuge tubes, and then subjected to a 5 min ultrasound bath. Then 0.2 mL heptane spiked with methyl-10-undecenoate (Sigma-Aldrich) as internal standard was added, followed by 1 min vortexing, and short centrifugation for phase separation. FA methyl esters were recovered in the heptane phase. Analyses were performed in a split-splitless injection mode on an AutoSystem XL Gas Chromatograph (Perkin-Elmer) equipped with a ZB-Wax capillary column (30 m x 0.25 mm x 0.25 μm; Phenomenex, France). Data were recorded and analyzed by TotalChrom Workstation (Perkin-Elmer). *S. aureus* FA peaks were detected between 12 and 32 min of elution, and identified with retention times of purified esterified FA standards.

### Detection of ACP products

Pellets from OD_600_=10 equivalent of exponential phase cultures were collected by centrifugation (8000 rpm for 5 min), and then washed twice with PBS (Phosphate Buffer Saline)-Triton X100 0.1%, and once with PBS. Pellets were re-suspended in 600 µL PBS supplemented with 6 µL antiprotease (100X), 50 µg/mL lysostaphin and 50 µg/mL DNAse and incubated with shaking for 25 min at 37°C. Protein concentrations were determined [47], and extracts (10 μg) were loaded onto non-denaturing gels containing 1M urea and 14% acrylamide gel as described [5, 7]. After electrophoresis, gels were subject to electroblotting for transfer to a PVDF membrane (Biorad, France) using a current of 25 V and 1.3 A on a power blotter semi-dry transfer system (Invitrogen, France) for 7 min. Primary anti-ACP antibody (Covalabs, France; [7]) was used at 1:500 dilution and secondary anti-rabbit IgG conjugated with peroxidase (Invitrogen, France) at a 1:5000 dilution. ACP-reactive bands were identified using the ECL kit (Perkin Elmer, France) and the fluorescent signal was recorded using ChemiDoc (BioRad, France).

### Phage transductions

Transductions of *fadM::*Tn transposon insertion from JE2 into Δ*plsX* suppressors *fadM2* and *fabF1* were performed as described using ϕ80 phage stock [48] [49]. The *fadM* transposon insertion [28] was transduced into JE2 Δ*plsX fadM2* suppressor strain 7.1 and into JE2 Δ*plsX fabF1* suppressor strain 7.8, giving rise respectively to Δ*plsX fadM::*Tn and Δ*plsX fabF1 fadM::*Tn, using erythromycin (5 µg/ml) as selection (**Table S4**). Insertions were confirmed by PCR.

### FASII bypass of WT and Δ*plsX* derivative strains

Anti-FASII bypass was evaluated as described [6]. Strains were grown overnight in liquid SerFA medium (SerFA, BHI containing 10 % newborn calf serum and an equimolar mixture of 83 µM each C14:0, C16:0, and C18:1 [Larodan, Sweden]). Cultures were then diluted to OD_600_=0.05 in SerFA containing or not 0.5 µg/ml of the anti-FASII AFN-1252 (here called AFN; [36]; MedChemExpress, France). Growth was monitored using the Spark plate reader (Tecan, France).

### Whole-genome sequencing data of S. aureus Δ*plsX* mutants and suppressors

Data is available in the Mendeley database: DOI: 10.17632/mzw2429w3m.1.

## Acknowledgements and funding

We are grateful to Kam Pou Han and Philippe Bouloc (Bouloc lab, I2BC, Orsay, France) for providing transducing phage and plasmid; Florence Dubois-Brissonet and Katia de Oliveira (Micalis, INRAE) for expert help and advice with CpG analyses; Candice Rigoulay for a simplified phage transduction protocol. We thank Micalis Institute colleagues Vincent Juillard for discussion of PlsX functions, and Philippe Gaudu, Jasmina Vidic, and team members for stimulating discussion and pertinent comments. We thank the PAPPSO proteomics facility (https://doi.org/10.15454/1.5572393176364355E12 ; supported by Paris-Saclay University, INRAE, AgroParisTech, the Ile-de-France Regional Council, and the IBiSA network) for an exploratory study that inspired β-lactam sensitivity testing.

This work was supported by the French Agence Nationale de la Recherche ANR (ANR-16-CE15-0013; AG), the Joint Programming Initiative on Antimicrobial Resistance (JPIAMR) under the umbrella of the ANR (ANR-22-AAMR-0007; AG), and the Fondation pour la Recherche Medicale (DBF20161136769; AG). PW was recipient of a Franco-Thai scholarship from Campus France and Khon Kaen University.

## Data availability

All relevant data are within the manuscript, supporting information files, and repositories. Any biological materials are available upon request.

## Author Contributions

All authors performed, analyzed, and interpreted experiments, and approved the manuscript. PW, JAM, and AG conceived the project, designed research, and wrote the paper.

## Supplementary Figures and Tables

**Fig. S1.**
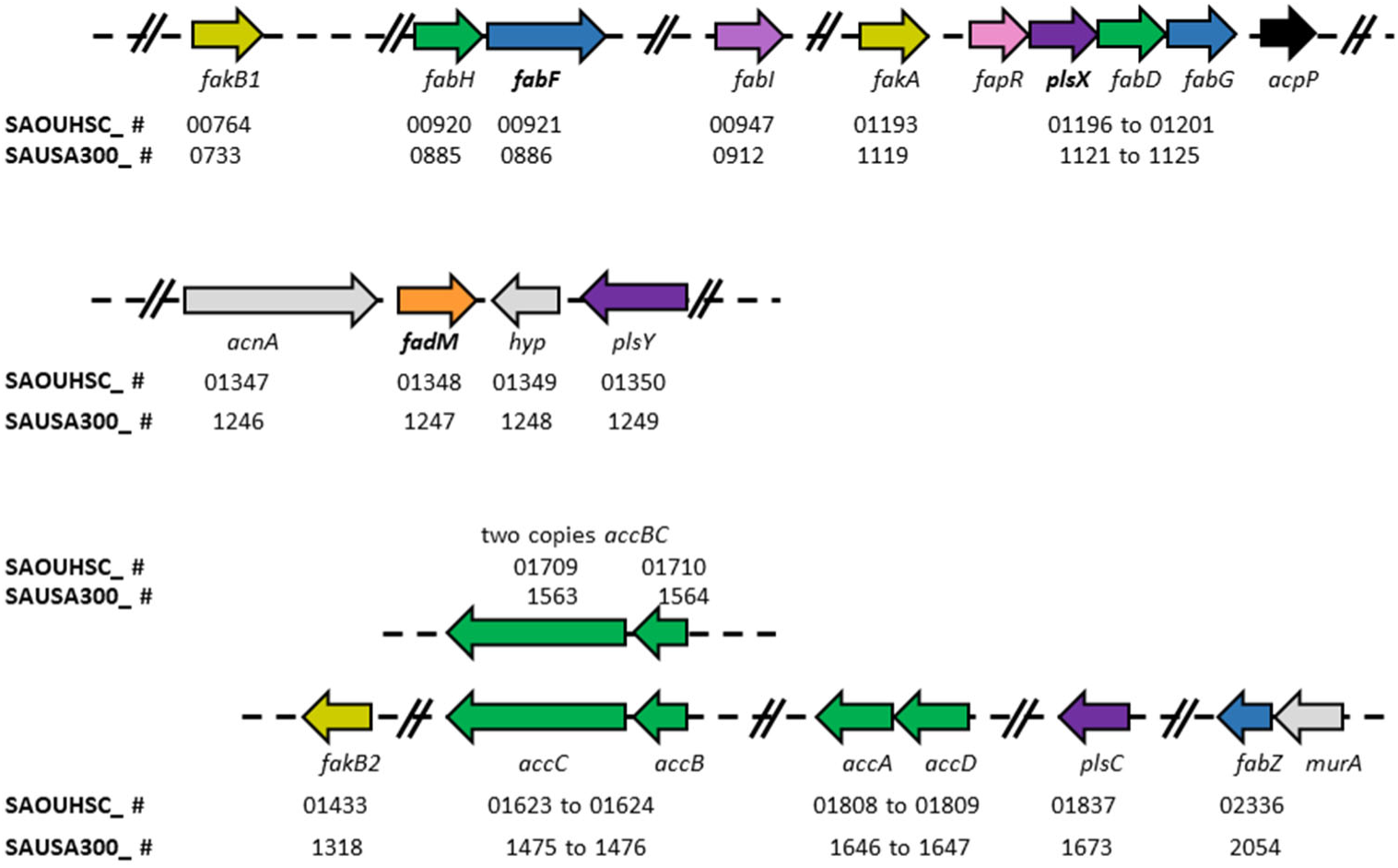
FASII and phospholipid gene organization in RN-R and JE2 strains. Locus tags of genes related to this study are presented. Dark green, FASII initiation; blue, FASII elongation; black, acyl carrier protein (*acpP*); purple, phospholipid synthesis; olive, FASII bypass fatty acid kinase; pink, regulatory; orange, FadM; grey, not directly related. Locus tags are given for USA300 FPR_3797 8325-4 as references.

**Fig. S2.**
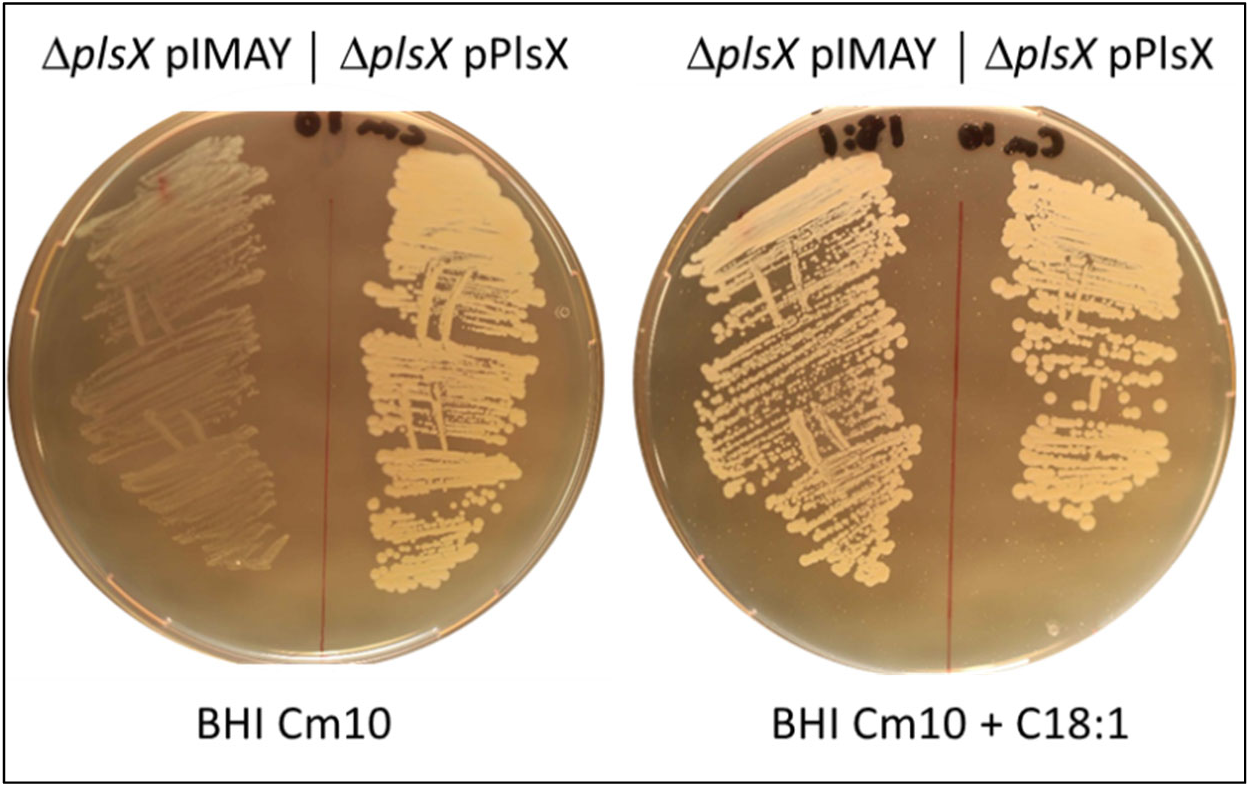
Complementation of Δ*plsX* by a plasmid-carried *plsX* gene. RN-R Δ*plsX* strains containing an empty plasmid carrying a chloramphenicol resistance cassette (pIMAY; left side of plates), or a plasmid carrying the *plsX* gene expressed from its native promoter on pIMAY (pPlsX; right side of plates), were streaked on BHI-chloramphenicol (Cm) 10 µg/ml solid medium, without or with 250 µM C18:1 to bypass the Δ*plsX* defect. Background growth on BHI (left) may be due to FA carryover from pre-cultures or to FA traces in medium. Plates were photographed after 48h growth at 30°C. The Δ*plsX* strain carrying the empty plasmid failed to grow in the absence of C18:1 supplementation, while the pPlsX-complemented strain grew in both media.

**Fig. S3.**
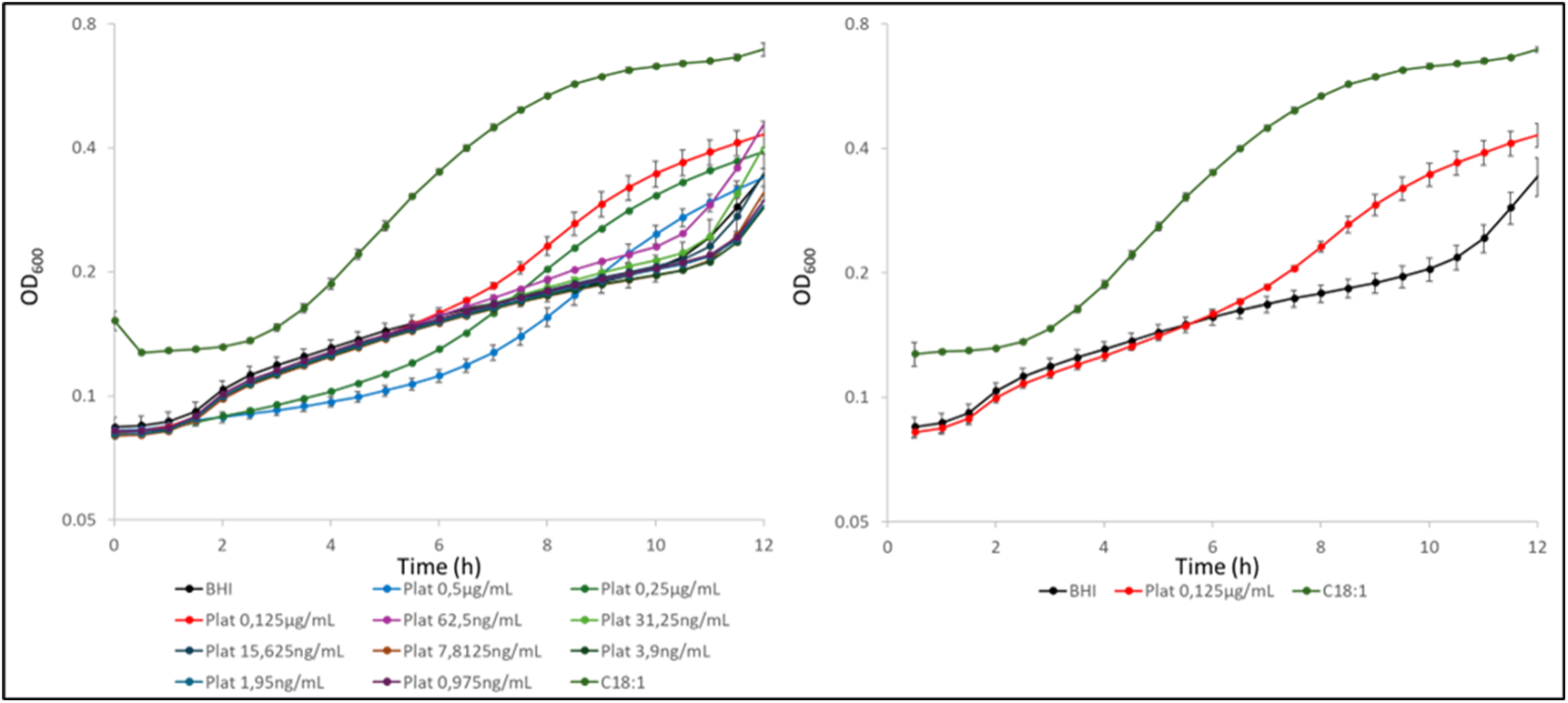
JE2Δ*plsX* growth stimulation by platensimycin in liquid culture. JE2 Δ*plsX* was grown overnight in BHI medium containing C18:1. After washing, the strain was resuspended to OD_600_ = 0.05 and used to inoculate BHI medium to which platensimycin was added at the concentrations indicated below graphs. Left, growth curves with the different platensimycin concentrations added to medium. Right, the optimal concentration leading to improved growth with platensimycin (red line) compared to growth in BHI medium (black line). The averages of three independent triplicates are shown. Differences are statistically significant (P=≤0.05) by 2-tailed bilateral T-test for all points comparing Δ*plsX* growth in BHI and BHI plus platensimycin 0.125 µg/ml) between 7 h and 12 h growth.

**Fig. S4.**
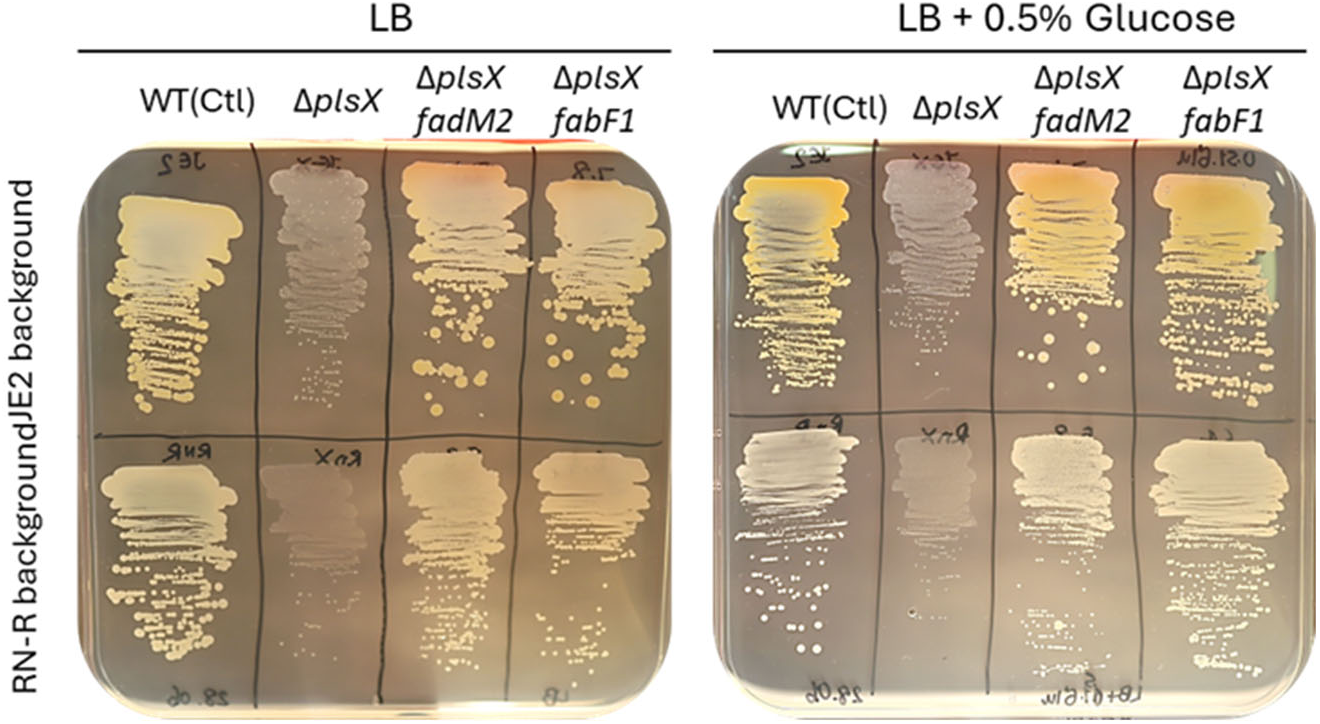
FadM^m^ acyl-CoA thioesterase activity is not needed for Δ*plsX* suppression. **A.** The FadM^m^ suppressor phenotype is not subject to glucose repression, suggesting that the Fad-mediated acyl-CoA synthesis degradation pathway (Fad) is not involved. Fad enzyme FadE converts FA to acyl-CoA, a FadM substrate [20]. The *fad* operon is subject to CcpA-glucose repression [25–28], suggesting that acyl-CoA pools would drop in glucose-containing medium. Strains precultured in LB and LB with C18:1 for Δ*plsX* (Materials and Methods) were streaked directly on solid LB medium without or with 0.5% glucose. Background growth on LB (left) may be due to FA carryover from pre-cultures or FA traces in medium. Upper and lower rows, strains are derived from JE2 and from RN-R, respectively. Results represent biological triplicates.

**Fig. S5.**
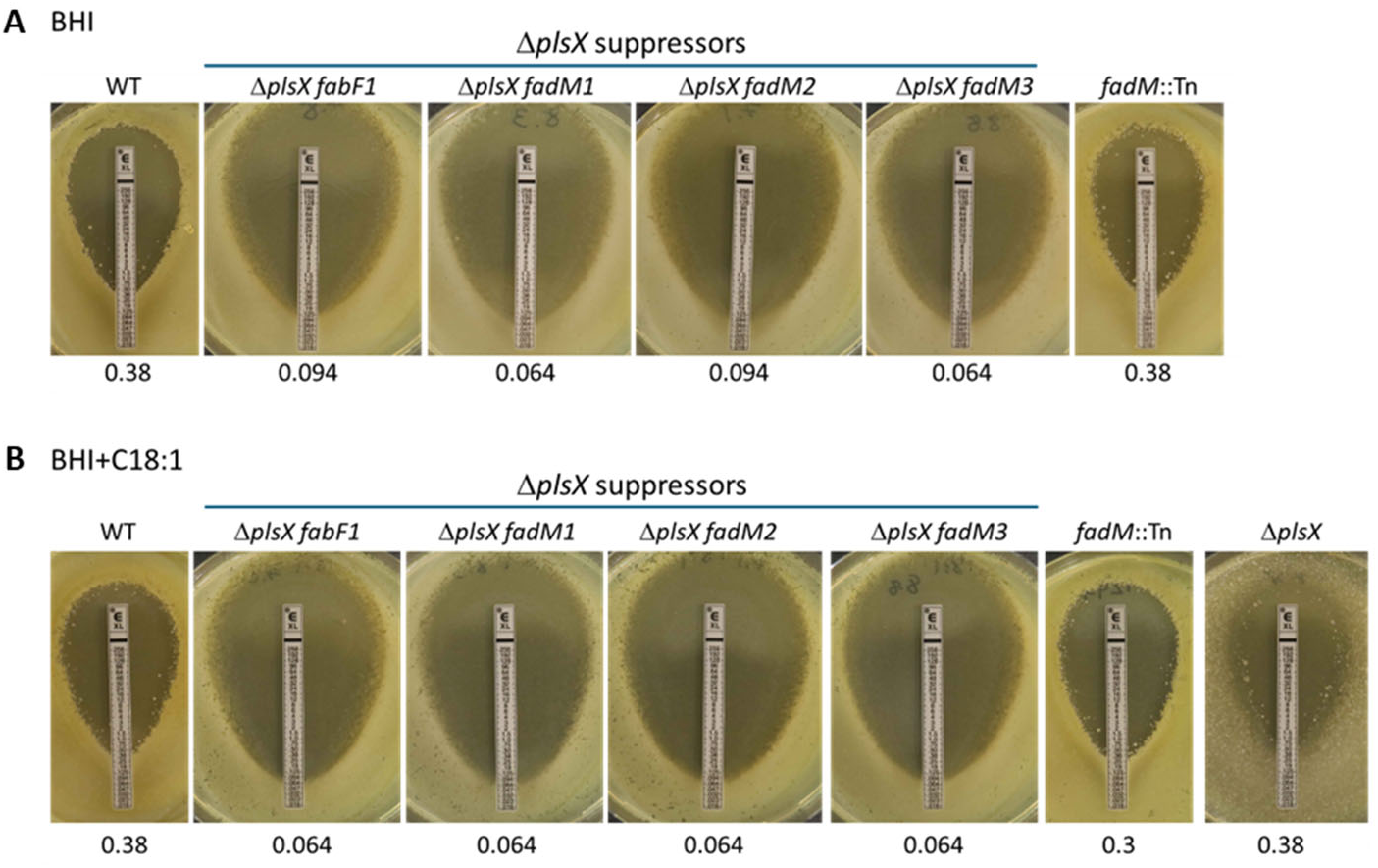
MRSA JE2 Δ*plsX* suppressor mutants are sensitized to the β-lactam antibiotic amoxicillin. The indicated JE2 and Δ*plsX* derivative strains were grown overnight in BHI medium without (**A**) or with 250 µM C18:1 (**B**). Cultures were resuspended to OD_600_ = 0.1 and 100 µl were spread on the cognate solid media. Amoxicillin Etest strips were placed on plates, followed by 48 h incubation at 37°C. The amoxicillin MICs are indicated in micrograms per milliliter below each photo.

**Fig. S6.**
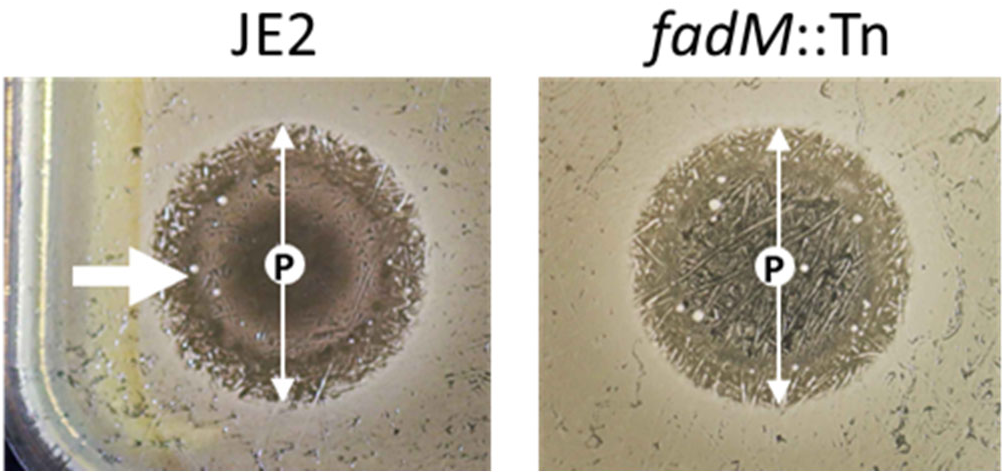
FadM is required for platensimycin-stimulated ring formation in the WT background. Overnight BHI cultures of JE2 and *fadM::*Tn (SAUSA300_1247) were washed in 0.9% NaCl, pellets were resuspended to OD_600_ = 0.1, and 250 µl was spread on 120 x 120 mM square petri plates containing solid BHI medium. Three µg platensimycin were spotted on plates (‘P’ in white circle; N=2). Plates were photographed at 48 h. Vertical arrow, inhibition zone; left arrow, growth ring surrounds the platensimycin deposit site in the WT, but not the *fadM::*Tn strain.

**Table S2.**
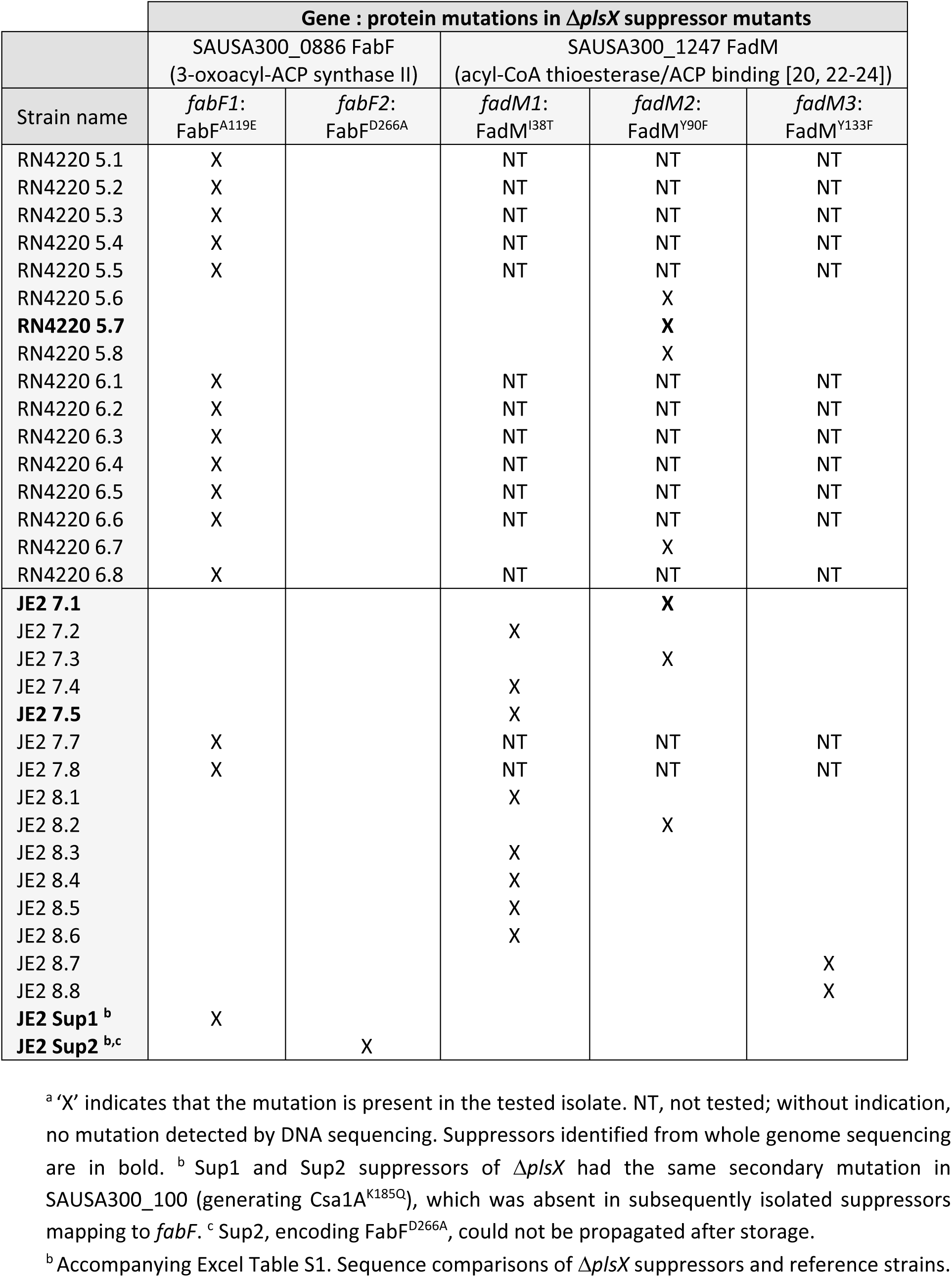
Δ*plsX* suppressors ^a,b^.

**Table S4.** Genetic materials.

| Strains |  |  |  |
| --- | --- | --- | --- |
| Host strain | Strain | Description | Reference |
| JE2<br>USA300_FPR3757 | JE2 (WT) | USA300_FPR3757 cured of its three native plasmids | [28] |
| | $\Delta plsX$ | In-frame *plsX* deletion leaving operon intact | This study |
| | $\Delta plsX$ suppressor in *fadM*; $\Delta plsX fadM1$ | *fadM1* encodes FadM<sup>I38T</sup> | This study |
| | $\Delta plsX$ suppressor in *fadM*; $\Delta plsX fadM2$ | *fadM2* encodes FadM<sup>Y90F</sup> | This study |
| | $\Delta plsX$ suppressor in *fadM*; $\Delta plsX fadM3$ | *fadM3* encodes FadM<sup>Y133F</sup> | This study |
| | $\Delta plsX$ suppressor in *fabF*; $\Delta plsX fabF1$ | *fabF1* encodes FabF<sup>A119E</sup> | This study |
|  | *fadM* Tn insertion | JE2 derivative containing an erythromycin-resistance-marked transposon (Tn) insertion in SAUSA300_1247 at position 1384915 (based on JE2 genome entry NZ_CP020619) | [28] |
| | $\Delta plsX fadM::Tn$ | $\Phi 80$ phage transduction of the SAUSA300_1247 Tn insertional mutant into the $\Delta plsX fadM2$ (7.1, **Table S2**) suppressor | This study |
| RN-R 4220 | RN-R 4220 (WT) | RN4220 derivative repaired for *fakB1* | [15] |
| | $\Delta plsX$ | In-frame deletion of *plsX* leaving operon intact | This study |
| | $\Delta plsX$ suppressor in *fadM*; $\Delta plsX fadM2$ | *fadM2* encodes FadM<sup>Y90F</sup> | This study |
| | $\Delta plsX$ suppressor in *fabF*; $\Delta plsX fabF1$ | *fabF1* encodes FabF<sup>A119E</sup> | This study |
| Plasmids |  |  |  |
|  | Plasmid | Description | Reference |
|  | pMAD | Thermosensitive plasmid used to generate double-crossover deletion events | [42, 43] |
|  | pMAD-*fapR-fabD* fragments | Used to generate *plsX* deletion | This study |
|  | pIMAY | Used for *plsX* cloning | [44, 57] |
| | pIMAY-*plsX* | Used for *plsX* complementation in $\Delta plsX$ | This study |
| Transducing phage |  |  |  |
|  | Phage | Description | Reference |
| | $\phi 80$ | Used to transduce erythromycin-resistance-marked Tn insertion of SAUSA300_1247 into $\Delta plsX fadM2$ and $\Delta plsX fabF1$ backgrounds | [58] |

**Table S5.**
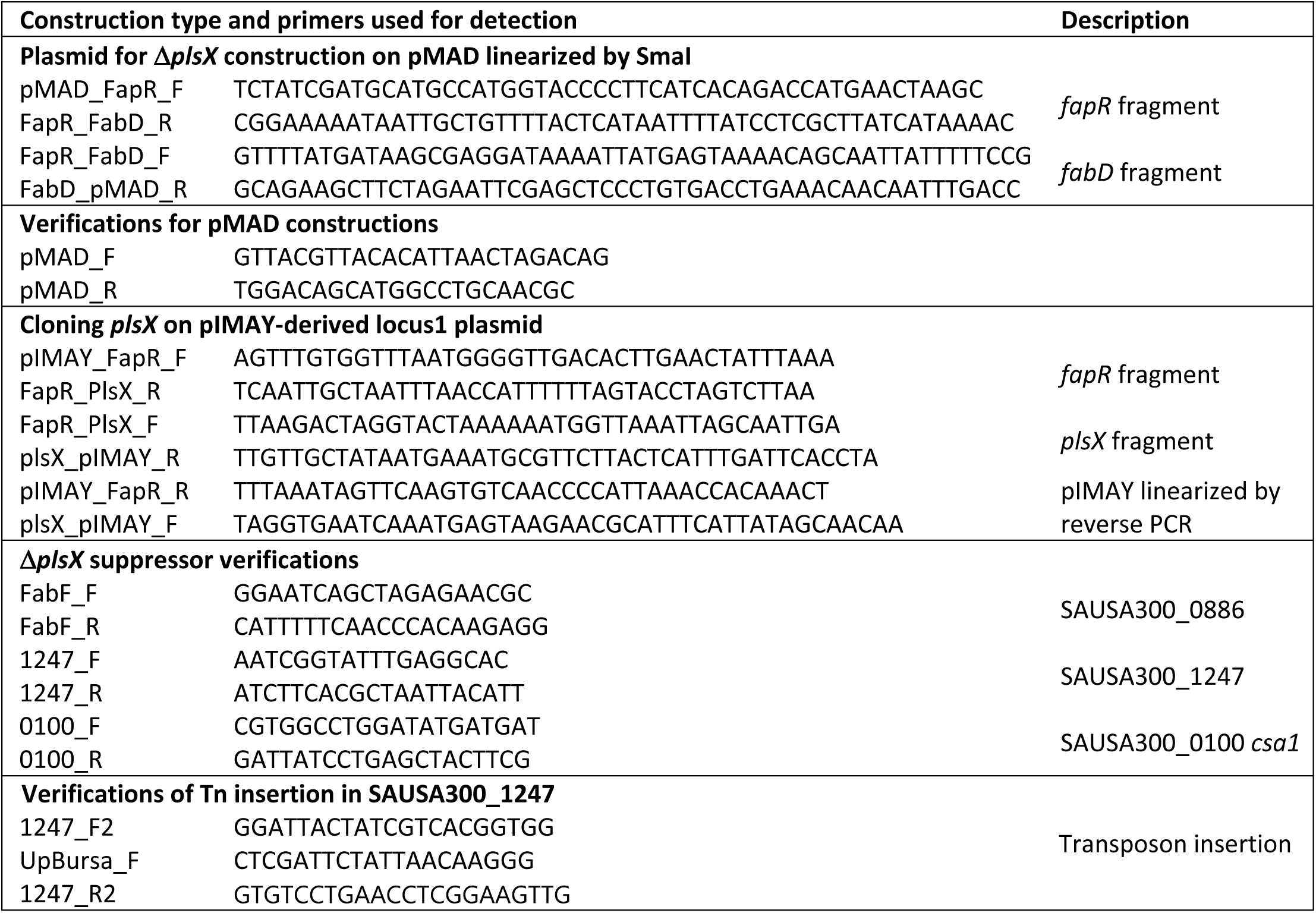
Oligonucleotides (5’ to 3’) used for genetic constructions and mutant detection.

